# The Colorado potato beetle gene expression atlas

**DOI:** 10.1101/2024.03.28.587222

**Authors:** Léonore Wilhelm, Yangzi Wang, Shuqing Xu

## Abstract

The Colorado potato beetle (CPB) is a major pest of potato crops that has evolved resistance to more than 50 pesticides. For decades, CPB has been a model species for research on insecticide resistance, insect physiology, diapause, reproduction and evolution. Yet, the research progress in CPB is constrained by the lack of comprehensive genomic and transcriptomic information. Here, building on the recently established chromosome-level genome assembly, we built a gene expression atlas of the CPB using the transcriptomes of 61 samples representing major organs and developmental stages. By using both short and long reads, we improved the genome annotation and identified 6,658 more genes that were missed in previous annotations. We then established a web portal allowing the search and visualization of the gene expression for the research community. The CPB atlas provides useful tools and comprehensive gene expression data, which will accelerate future research in both pest control and insect biology fields.

## Background & Summary

The Colorado potato beetle (CPB) is a devastating pest originating from North America that has spread in Europe and Asia in the 20^th^ century. It causes substantial damage to solanaceous plants, in particular potatoes, as one CPB can consume around 40 cm^2^ leaf area during the larval stages and 100 cm^2^ leaf area every 10 days during its adult life ^1^. The CPB has a remarkable resistance to pesticides. More than 50 active compounds proved to be ineffective against it ([Mota-Sanchez and Wise 2017 Arthropod Pesticide Resistance Database](https://www.pesticideresistance.org/display.php?page=species&arId=141)). Studies have shown that the resistances to many insecticides evolved rapidly in response to the applications of insecticides and showed large geographical variations ^2–6^. This is largely due to high genetic variation in the population ^7^, likely as the results of CPB’s high fecundity. A mated female CPB can produce 25 eggs per day on average ^8^ and up to 724 eggs in 30 days ^9^, generating a large pool of individuals upon which selection can act. Adaptation to pesticides appears to be polygenic, involving genes related to detoxification, cuticle composition, and neuronal receptors ^7,10^.

The CPB has been the object of extended studies, most of which are related to understanding the mechanisms of pesticide resistance. Agronomic studies on the CPB include bioassays with potential pesticides ^11,12^, use of biocontrol agents ^13^, and RNA interference ^14–17^. Recent studies have focused on finding alleles linked with resistance ^18,19^ and gene expression linked with resistance ^20^. The CPB has also been used as a model to understand insect development and physiology ^21–23^, insect diapause ^24–27^, and more recently insect lipid metabolism and calcium signaling ^28,29^ as well as behavior ^30^.

Despite the abundance of research topics addressed using the CPB, the research in CPB is currently constrained by the lack of accessible genomic and transcriptomic data. Until recently, the chromosome-level reference genome has been released and several studies on population genomics of CPB have been carried out ^31,32^. Yet, we currently have no publicly available gene expression data among tissues and developmental stages, which is vital for understanding the genetic mechanisms of most phenotypic traits in CPB. Here, we sequenced the transcriptomes of 12 tissues in adults (five from female, seven from male), five tissues in last instar larvae, as well as the entire body of each larval stage, and eggs. Using these data, we further improved the genome annotation and established a gene expression atlas of CPB. The updated genome annotation contains 34,350 genes, with a BUSCO score of 96.2%, which is 3% higher than previous annotation. To provide an easy access to the expression data, we established a web-based portal allowing users to search, download and visualize the expression of genes among organs or developmental stages. The CPB atlas can be accessed via https://cpb-atlas.uni-mainz.de/

## Methods

The insects were maintained in a climate chamber with 24°C, 70% humidity and a photoperiod of 16 h light and 8 h dark per day. The E06 strain, which was originally collected from Spain in 2012, was used for all samples^33^. The experimental design comprises the following steps (Figure 1): dissection of the specimens, RNA extraction, RNA sequencing, data processing, and generation of the functional annotations.

**Figure 1.**
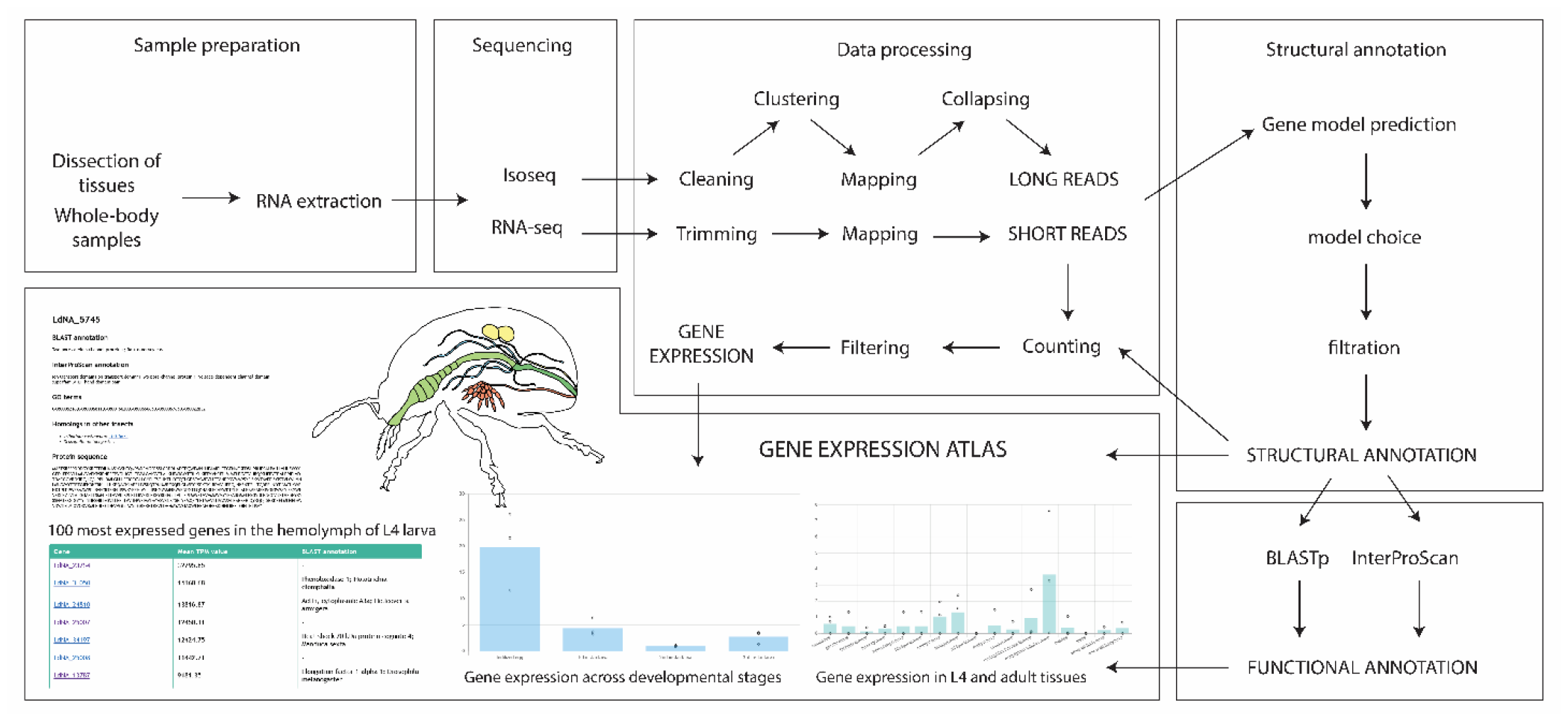
Graphical representation of the CPB atlas workflow. The samples were first prepared (either dissected or directly frozen), RNA was extracted, then sequenced. The data from RNA-seq and Isoseq were processed separately. Both were used to construct the structural annotation i.e. position of the genes and their exons and CDS on the genome assembly. The structural annotation was employed later to build the functional annotation i.e. function of each gene, by a search with BLASTp and InterProScan separately. The short reads were mapped to the structural annotation to obtain an expression count for the genes. The resulting gene expression were displayed on a web portal in the form of bar plots.

### Dissection and RNA extraction

The CPB atlas includes transcriptomes of tissues from fourth instar larvae (L4) and adults, as well as the whole body from different stages, from egg to third instar larva (figure 2). The tissues were dissected in PBS under a stereoscopic microscope. Dissection time was limited to 10 min to avoid RNA degradation and the tissues were subsequently snap-frozen in liquid nitrogen and stored in a -80°C freezer. The whole-body samples were similarly snap-frozen before being stored to ensure identical processing of the entire dataset. The eggs were pooled into groups of five. Three L1, L2, and L3 larvae were pooled to create each sample, respectively. The L4 larvae and adult tissues are from single individuals. The L4 larvae were dissected on the 19^th^ day of development from egg laying. Adults were kept in individual petri dishes from emergence and dissected seven days later. RNA was extracted using the RNeasy Mini kit from Qiagen (Venlo, The Netherlands), following the manufacturer’s protocol. Lysis was performed with micro pestles. To obtain the full-length transcript of all the genes using an iso-seq approach, we pooled 200 ng RNA from one replicate of each sample type (except two due to their low RNA abundance; Supplementary table 1). The 61 RNA-seq samples and one iso-seq sample were sequenced using Illumina NovoSeq and PacBio Sequel sequencers at Novogene (Cambridge, UK), respectively. We used three replicates for each sample except the white fat body and the yellow fat body of the L4 larva, for which two samples were used.

**Figure 2.**
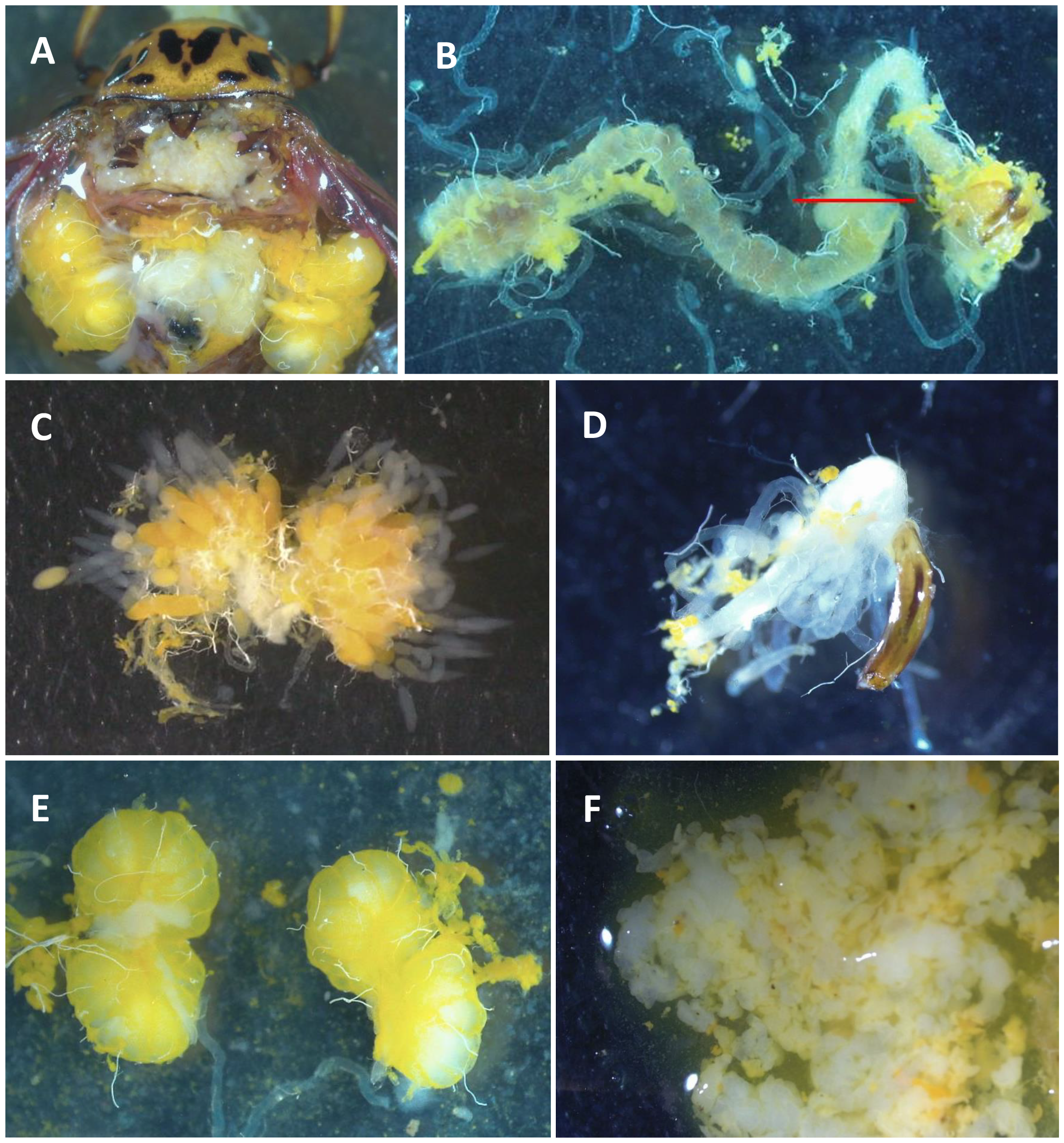
Dissection of tissues: A. Male incised dorsally. B. Midgut, hindgut and malpighian tubules; C. Ovaries; D. Aedeagus and genital ducts; E. Testis; F. Fat body (white and yellow) of a fourth instar

### Iso-seq data processing

Iso-seq data were processed using the command-line tools from PacBio’s SMRT Link software (available at https://www.pacb.com/). First, consensus reads were generated with the function *css*. We demultiplexed reads with *lima* and then removed polyA tails with *isoseq3 refine*. The reads were clustered with *isoseq3 cluster*. The Full-Length Non-Concatemer (FLNC) reads obtained were mapped to the genome ^31^ using pbmm2, the pacbio wrapper for minimap2 ^34^. Finally, they were collapsed using Cupcake (v0.1.4, https://github.com/Magdoll/cDNA_Cupcake) to produce non-redundant full-length transcripts.

### RNA-seq data processing

Trimmomatic was used to remove the Illumina adapters, drop the reads with a low quality or a short length (SLIDINGWINDOW:4:15 MINLEN:36) and remove the first ten bases of the reads (HEADCROP:10). To assess the quality both before and after this filtering process, we utilized FastQC (v0.11.9, https://www.bioinformatics.babraham.ac.uk/projects/fastqc/). The reads were mapped to a chromosome-level genome assembly ^31^ using STAR ^35^ (version 2.7.8a). FeatureCounts (Subread 2.0.5) was used to count the reads mapped to the annotated genes, with the parameters “-p –countReadPairs” which indicates that the reads are paired.

The genes that were not expressed were filtered out based on their transcript per million (TPM) values. We kept only the transcript having a TPM value above one in at least two samples (script available on the Github). After filtration, 15,578 genes out of 34,350 were kept in the gene expression atlas.

### Protein-coding gene annotation

For the prediction of protein-coding genes, we employed a modified BRAKER^36–40^ pipeline (https://github.com/Gaius-Augustus/BRAKER/blob/master/docs/long_reads/long_read_protocol.md). In brief, this approach integrates gene models predicted based on both short-read RNA-seq transcriptome (BRAKER1 method ^41–46^) and protein homologs from *Drosophila melanogaster* and *Tribolium castaneum* (BRAKER2 method ^39,42,43,47–51^). Subsequently, TSEBRA ^40^ was used to compare these predictions against full-length transcripts obtained from Iso-seq data, thereby determining the most accurate gene models.

For running BRAKER1, we provided with paired-end RNA-seq reads from 17 different tissue types (Supplementary table 2) and the repeat soft-masked CPB reference genome ^31^. The short reads facilitated the automatic training of GeneMark-ET by using the spliced alignment data to assist gene model predictions by AUGUSTUS ^52^. For BRAKER2, a similar process was conducted for the automatic training of GeneMark-EP+ ^51^. However, in this step, BRAKER2 employed information on protein-coding exon boundaries derived from the alignment of homologous protein sequences from *Drosophila melanogaster* and *Tribolium castaneum* (downloaded from UniProt ^53^) to assist AUGUSTUS in gene prediction. We used GeneMarkS-T ^54^ to identify the protein-coding regions within each full-length transcript. We then subsequently merged gene models from BRAKER1 and BRAKER2 and compared against the Iso-seq full-length transcripts using TSEBRA. We retained only the longest isoform for each gene model. Using AGAT (v0.9.0, https://github.com/NBISweden/AGAT) we removed genes that are shorter than 100 bp and single-exon genes that lacks complete start or stop codon. Unique gene models that were present in the previous annotation version ^31^ and had no overlap with any models predicted in the current version were revived and incorporated into the new annotation. For the functional annotations, protein sequences were aligned to the UniProtKB^20^ database using blastp (BLAST+ v2.12.0) ^43^ with “-evalue 1e-6 - max_hsps 1 -max_target_seqs 1 -outfmt 6”. The results were further processed using InterProScan (v5.63-95.0) ^55^ with the options “-goterms -iprlookup”.

## Data Records

The raw sequences generated for this paper have been deposited on NCBI under the bioproject accession number PRJNA1067435.

## Technical validation

The sample integrity, purity, and quantification were verified using an Agilent 5400 device. Samples were visualized on an NMDS and a pairwise heatmap to detect potential outliers using the log2(TPM+1) values (Supplementary figures 1 and 2). Two outliers were detected, a yellow fat body and a female hindgut. As this is likely due to the contaminations during dissection and sample collection steps, we removed these two samples from the CPB atlas. Both visualization methods were applied again and show that replicates cluster together (Figures 3 and 4).

**Figure 3.**
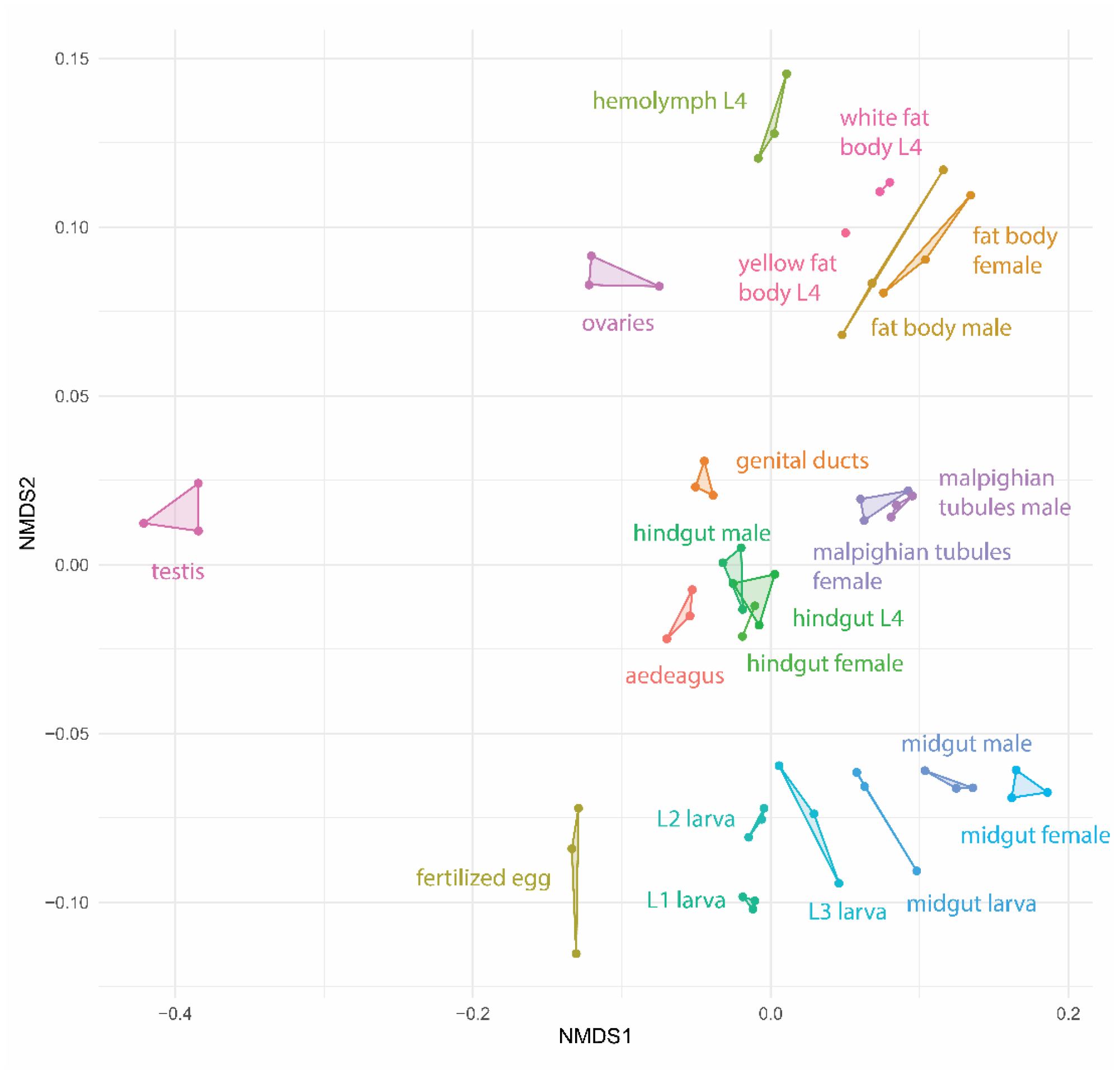
Non-metric multidimensional scaling (NMDS) of the samples based on log2(TPM+1) values. Biological replicates are connected.

**Figure 4.**
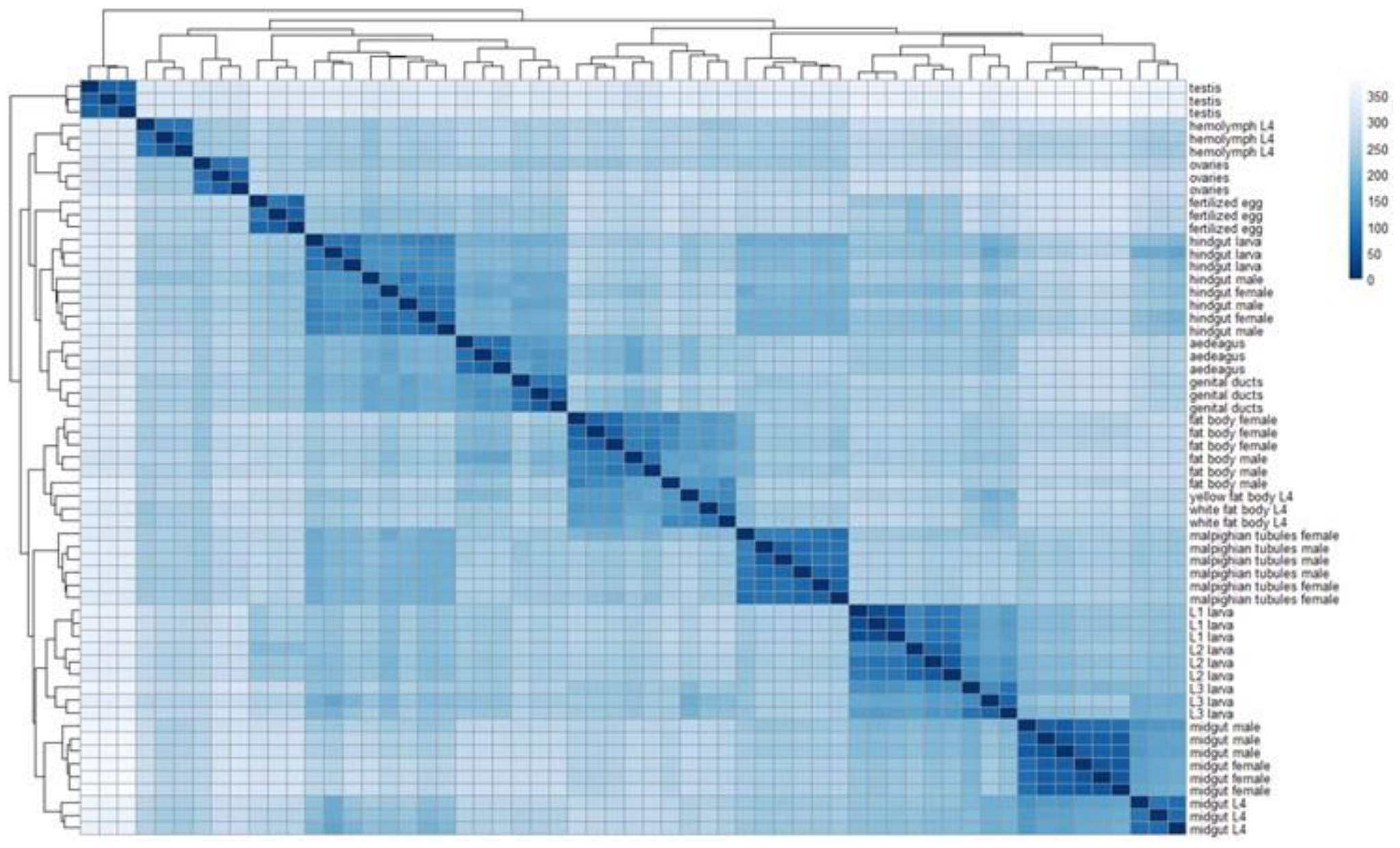
Heatmap showing the pairwise comparisons of log2(TPM) values across the 59 samples of the gene expression atlas. All the tissues are correctly clustering together showing an absence of contaminations from other tissues. Some tissues also segregate by developmental stage (e.g. midgut of L4 larvae vs adults) showing a modification of the transcriptome through age. On the contrary there are no clear separation of the tissues between the sexes in adult, showing the absence of sex-specific transcriptome for tissues common to both sexes.

## Code availability

The scripts used to process short reads are available at the adress: https://github.com/Xu-lab-Evolution/CPB_gene_expression_atlas. The genome annotation has been generated with the method described at: https://github.com/Xu-lab-Evolution/Waterlily_aphid_genome_project

## Acknowledgment

We acknowledge Cansu Doğan for her advice on identifying and dissecting the CBP tissues, Ursula Martiné for her considerable help in the RNA extraction, Christina Sternara Tsiara and Pablo Duchen for creating the heatmaps displayed on the web application, Alex Garel for deploying the web application. This project was funded by the Deutsche Forschungsgemeinschaft (DFG, German Research Foundation) – GRK2526 (Project number 407023052).

## Author contributions

Conceived and designed the experiments: S.X., L.W. Performed the experiments: L.W. Analyzed the data: L.W., Y.W. Built the web application: L.W., Wrote the paper: L.W., Y.W. Substantially revised the paper: S.X. Supervision and grant acquisition: S.X.

---

**Table 1.** Comparison of the functional annotation from Yan et al. with the annotation from this paper.

| Annotation version | Data used | Gene number | Busco score |
| --- | --- | --- | --- |
| Yan et al 2023 Sci Data | • 13 RNA-seq samples of whole-body of adults and 1<sup>st</sup> instar larvae<br>• Protein homologs of all insects downloaded from OrthoDB | 27692 | C:93.5%[S:86.0%,D:7.5%]<br>,F:1.0%,M:5.5%,n:1367 |
| This paper | • 17 RNA-seq samples from different tissues and developmental stages (Supplementary table 2)<br>• Protein homologs from *D. melanogaster* and *T. castaneum* downloaded from Uniprot<br>• Long reads from 21 sample types<br>• Genes annotated in Yan et al. but absent in our current annotation were added to the annotation. | 34350 | C:96.2%[S:88.1%,D:8.1%]<br>,F:0.9%,M:2.9%,n:1367 |

