## Supplementary files for "The Colorado potato beetle gene expression atlas"

**Supplementary table 1. Samples for which an extract have been collected and pooled for long read sequencing.**

| RNA-seq ID | Tissue types |
|---|---|
| A1 | aedeagus |
| D2 | genital ducts |
| FbF6 | fat body female |
| FBM7 | fat body male |
| FE2 | fertilized egg |
| HF5 | hindgut female |
| HL1 | hindgut larva |
| HM5 | hindgut male |
| HmK2 | hemolymph |
| ML4 | midgut larva |
| MM5 | midgut male |
| MTF1 | malpighian tubule female |
| MTM3 | malpighian tubule male |
| O4 | ovaries |
| T2 | testis |
| L1_1 | whole-body first instar larva |
| L2_1 | whole-body second instar larva |
| L3_2 | whole-body third instar larva |

**Supplementary table 2. Samples provided to BRAKER to build the structural annotation.**

| RNA-seq ID | Tissue types |
|---|---|
| A2 | aedeagus |
| D1 | genital ducts |
| FbF2 | fat body female |
| FBM1 | fat body male |
| FE1 | fertilized egg |
| HF1 | hindgut female |
| HL1 | hindgut larva |
| HM3 | hindgut male |
| HmK3 | hemolymph |
| ML3 | midgut larva |
| MM2 | midgut male |
| MTF1 | malpighian tubule female |
| MTM3 | malpighian tubule male |
| O1 | ovaries |
| T1 | testis |
| WFB1 | white fat body larva |
| YFB2 | yellow fat body larva |

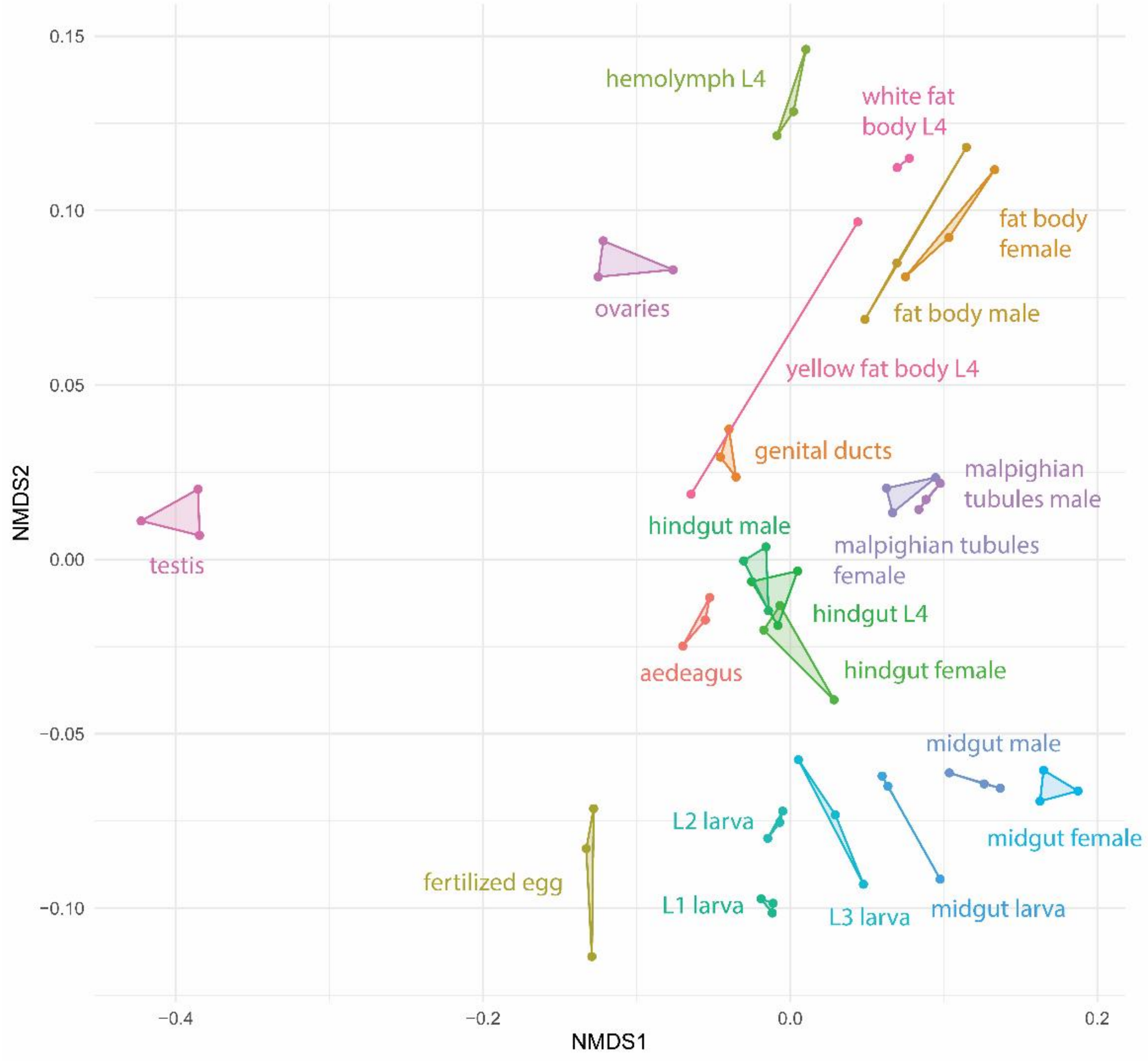


**Supplementary figure 1.** Non-metric multidimensional scaling (NMDS) of the samples based on log2(TPM+1) values. Biological replicates are connected. A yellow fat body of L4 appear clearly as an outlier, it has been removed from the dataset.

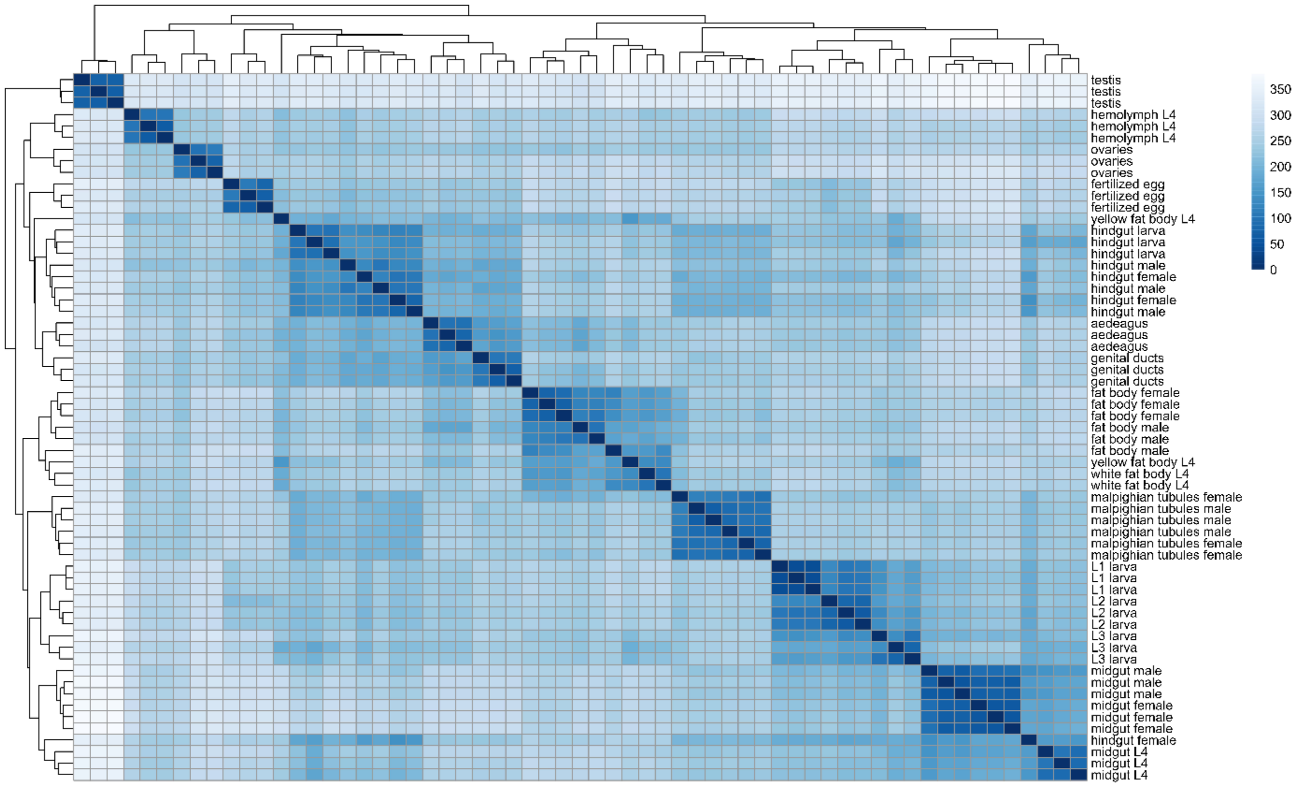


**Supplementary figure 2.** Heatmap showing the pairwise comparisons of log2(TPM) values across the 61 sequenced samples. A yellow fat body and a hindgut of female don't cluster with the replicates of the same type. Thy have been removed from the dataset.
